# Does a history of co-occurrence predict plant performance, community productivity, or invasion resistance?

**DOI:** 10.1101/2022.09.20.508783

**Authors:** Alison C. Agneray, Matthew L. Forister, Thomas L. Parchman, Elizabeth A. Leger

**Affiliations:** Graduate Program in Ecology, Evolution, and Conservation Biology, Department of Biology, University of Nevada, Reno, 1664 N. Virginia Street, Mail Stop 314, Reno, NV 89557; Bureau of Land Management Nevada State Office, 1340 Financial Blvd., Reno, NV 89502

**Keywords:** allopatry, community evolution, complementarity, co-selection, invasion resistance, niche differentiation, plant productivity, species interactions, sympatry

## Abstract

A history of species co-occurrence in plant communities is hypothesized to lead to greater niche differentiation, more efficient resource partitioning, and more productive, resistant communities as a result of evolution in response to biotic interactions. We asked if individual species or community responses differed when communities were founded with species sharing a history of co-occurrence (sympatric) or with species originating from different locations (allopatric). Using shrub, grass, and forb species from six locations in the western Great Basin, USA, we compared establishment, productivity, reproduction, phenology, and resistance to invaders for experimental communities with either sympatric or allopatric associations. Each community type was planted with six taxa in outdoor mesocosms, measured over three growing seasons, and invaded with the annual grass *Bromus tectorum* in the final season. For most populations, the allopatric or sympatric status of neighbors was not important. However, in some cases it was beneficial for some species from some locations to be planted with allopatric neighbors, while others benefited from sympatric neighbors, and some of these responses had large effects. For instance, the *Elymus* population that benefited the most from allopatry grew 50% larger with allopatric neighbors than in single origin mesocosms. This response affected invasion resistance, as *B. tectorum* biomass was strongly affected by productivity and phenology of *Elymus* spp., as well as *Poa secunda*. Our results demonstrate that while community composition can in some cases affect plant performance in semi-arid plant communities, assembling sympatric communities is not sufficient to ensure high ecosystem services. Instead, we observed a potential interaction between sampling effects and evolutionary history that can create invasion resistant allopatric communities.

## Introduction

Plant species that are adapted to similar environmental conditions can co-occur long-term if there are mechanisms facilitating coexistence (Chesson, 2000; Keddy, 1992; Levine et al., 2017). Niche differentiation is one process that allows coexisting species to partly escape competition for resources such as water, nutrients, and light (Godoy et al., 2020; Silvertown, 2004). Character displacement as a result of niche differentiation is a complex outcome of eco-evolutionary processes which can vary across time, spatial scales, and ecosystems (Eisen & Geber, 2018; Pearman et al., 2008; Thorpe et al., 2011). Though not ubiquitous, niche differentiation has commonly evolved in temperate plant species, resulting in plant communities with a mixture of rooting depths, varying phenology, alternative life histories, and differing leaf morphology (Bakker et al., 2021; Hector et al., 2010; Kulmatiski et al., 2020; Silvertown, 2004). This niche differentiation can lead to complementarity and facilitation between interacting plant neighbors, resulting in productive plant communities with a variety of species that partition available resources across time and space (Camarretta et al., 2020; Grady et al., 2017; van Moorsel et al., 2018).

In addition to the many observations of niche differentiation among interacting species, geographic variation in species interactions also commonly leads to trait differentiation among populations of the same species (Fernandes et al., 2019; Thompson, 2005; Thorpe et al., 2011). Variation in local interactions can generate differences in functional traits among populations of the same species, and lead to differences in niche overlap among species with a history of sympatry (shared evolutionary history at a single growing site) versus allopatry (divergent histories sourced from different sites; Eisen & Geber, 2018; Kooyers et al., 2017; Thorpe et al., 2011; Zuppinger-Dingley et al., 2014). Most work describing the impacts of evolutionary history on niche differentiation in plant populations has been performed on pairs of closely related species (Eisen & Geber, 2018; Kooyers et al., 2017), experimentally altered populations in grasslands (Chanway et al., 1988; van Moorsel et al., 2018; Zuppinger-Dingley et al., 2014), small numbers of species within the same functional groups (Camarretta et al., 2020; Grady et al., 2017), or through theoretical models (Aubree et al., 2020). There is evidence that a history of sympatry can positively affect individual plant performance, though in experiments varying the number of interacting species, the strongest evidence has been found with fewer interacting species from simulated or experimentally created communities (Aubree et al., 2020; van Moorsel et al., 2018). This raises the question of whether these dynamics can be observed in natural systems or with a greater number of species and functional groups.

Beyond effects at the species level, there has long been interest in emergent properties that result from plant interactions, especially the question of whether there are community-level outcomes that differ when co-occurring species have a history of coevolution (Aarssen 1983; Chanway et al., 1988; Whitman et al., 2020). For example, the traits within populations that are shaped from a history of interaction with sympatric species could lead to communities with greater primary productivity, resilience in the face of disturbance, and resistance to invasion due to increased complementarity and facilitation, if efficient resource use leads to overperformance in a coevolved community (Germain et al., 2016; van Moorsel et al., 2021; Zuppinger-Dingley et al., 2014). The best-studied of such responses is overyielding, wherein plant communities demonstrate enhanced primary productivity when they have a history of co-occurrence (Chanway et al., 1988; Grady et al., 2017; van Moorsel et al., 2018). In some experiments, mechanisms for greater productivity have been attributed to asynchronous phenology and less root overlap allowing plants to fully utilize available resources and increase aboveground biomass (Zuppinger-Dingley et al., 2014). However, one critique of studies comparing communities with sympatric vs. allopatric histories is that community-level effects are often assumed from observations of trait differences in sympatric populations, rather than measured directly (Germain et al., 2018).

Even in the absence of coevolutionary dynamics, there may be reasons why sympatric communities behave differently than assemblages of species with allopatric origins. If communities are assembled as a result of coincidental dispersal histories or ecological sorting, they may share evolutionary responses to the same environment (Gleason, 1926; Hubbell, 2001; Keddy, 1992). Thus, contrary to predictions about niche partitioning in interacting species, a shared environment could lead to shared characteristics and reduced niche differentiation in sympatric populations. For example, in stressful arid environments, greater root allocation and earlier emergence are often favorable and evolve in multiple species from different functional groups (Agneray et al., 2022; Baughman et al., 2019). Because the relative importance of plant competition decreases when environmental stress is high (Grime, 1977; Martínez-Blancas & Martorell, 2020), one might expect less evidence for character displacement among interacting species from low resource environments (Eisen & Geber, 2018; Kooyers et al., 2017; Thompson, 2005). Moreover, niche differentiation and complementarity are not required mechanisms for observing differences in community function: many studies have observed that emergent properties like productivity can be driven by sampling or dominance effects. For example, in experimentally assembled communities, certain populations or species with above average biomass can dominate the system and have outsized impacts on the resulting community biomass (Godoy et al., 2020; Mahaut et al., 2020).

Here, we evaluated the impact of occurrence history on interactions between multiple functional groups using wild-collected seeds of perennial grasses, forbs, and shrubs from the Great Basin US, a semi-arid system. This region encompasses a vast expanse of contiguous, undeveloped land, but disturbance and invasion, primarily from introduced annual grasses, make this area of high interest for conservation and restoration, including large-scale seeding to found communities of wild plants (Davies et al., 2011) We tested whether sympatric or allopatric histories affected species and community-level outcomes by growing the same set of six co-occurring species from six natural communities in experimental mesocosm communities for three growing seasons. Each community was assembled from either sympatric neighbors (six communities), or in six randomly assigned allopatric communities, creating 12 different communities that differed in interaction history but contained the same six species. We asked three questions: 1) do plants perform differently when grown with sympatric or allopatric neighbors, and do responses differ among species or populations? 2) do community-level responses (total productivity, total survival, total flower production, or invasive suppression) differ among allopatric or sympatric treatments? and 3) can we identify factors that can predict differences in invasion resistance among experimentally assembled communities? These questions are important for understanding how wild plant communities form and function, and in addition, are critical for guiding ecological restoration, which commonly assembles new communities using seeds from different locations (Kettenring et al., 2014).

On the one hand, based on theory and work from many temperate communities, one might expect that individual plants would perform better (grow larger, produce more flowers, and have higher survival) when grown with neighbors having a sympatric history of co-occurrence. As a result of this increased individual performance, sympatric mixes would be expected to have above average performance at a community level relative to allopatric communities due to their potentially differentiated characteristics. In an applied context, a consistent, positive effect of planting sympatric populations could lead to improved restoration outcomes and increased ecosystem function. On the other hand, semi-arid systems are underrepresented in the experimental community literature, and an alternative set of expectations include a reduced importance for sympatric histories and competition (Callaway et al., 2002), and a greater role for environmental filtering in shaping plant characteristics in resource-poor environments. Finally, we hypothesized that the performance of perennial grass species would have an outsized effect on invasion resistance based on their functional group. We expected that perennial grasses would compete more directly with the invasive, annual grass, due to their similar phenology and previous work showing the potential for strong interactions, including natural selection in response to invasion (Leger & Goergen, 2017).

## Methods

### Species selection and seed collection

We focused our work within the sagebrush-dominated regions of the western Great Basin. This semi-arid region is characterized by cold winters and hot, dry summers with highly variable precipitation that falls primarily in the winter as both rain and snow (West, 1983), and water is a primary resource limiting plant growth (Donovan & Ehleringer, 1994). Plant communities in the Great Basin are mixtures of native shrubs, perennial grasses, and perennial and annual forbs, along with a robust component of introduced and invasive annual grasses and forbs (West, 1983). In particular, *Bromus tectorum* L., or cheatgrass, is an annual invasive grass that is now ubiquitous in the sagebrush steppe and is highly competitive with native plants (Melgoza et al., 1990). Most native and introduced grasses germinate after the first fall rainstorms following a period of dormancy in the hottest summer months. Forbs are more variable, germinating in the fall, winter, or spring depending on temperatures and life history strategy (Barga et al., 2018), while shrubs typically germinate in the late winter and early spring after a period of cold stratification (Bonner & Karrfalt, 2008).

Target species were selected after conducting plant surveys in 21 locations in western Nevada, eastern California, and southeastern Oregon, with the goal of identifying the most commonly co-occurring plant community dominants. We selected six of the most common native plant species representing a variety of life forms. Species included three perennial grasses (*Elymus* spp. L., *Achnatherum thurberianum* (Piper) Barkworth, *Poa secunda* J. Presl), two shrubs (*Artemisia tridentata* Nutt. and *Ericameria nauseosa* (Pall. ex Pursh) G.L. Nesom & Baird), and one annual/biennial forb (*Chaenactis douglasii* (Hook.) Hook. & Arn.). We included sites where *Elymus elymoides* (Raf.) Swezey and *E. multisetus* M.E. Jones co-occur and likely hybridize, given their shared distribution in the western Great Basin (Barkworth et al., 2007). For simplicity, we refer to *Elymus* taxa jointly as a “species”, even though some *Elymus* collections may include two hybridizing species.

We identified six collection sites with similar climatic conditions where all focal species occurred (Figure S1). Sites were primarily mid-elevation, sagebrush steppe communities, with average annual precipitation between 236 and 383 mm and elevation ranging from 1395-2055 m (PRISM Climate Group, 2004). We bulk-collected mature seeds from a minimum of 50 individual plants for each species from each location between 1 June and 15 December of 2015.

### Experimental mesocosms

We prepared 90 mesocosms (Figure S2; 200L) outside of the Valley Road Greenhouse Complex at the University of Nevada, Reno (39.537924, −119.804757). Each mesocosm was filled with locally sourced topsoil to 0.9m depth (soil surface area = 0.25m^2^, soil volume =0.3m^3^). Seeds were examined on a light table to ensure seed fill, and planted in November 2016, following the same orientation and planting design in each mesocosm (Figure S2). Each mesocosm included two replicates of *Elymus* spp. and *A. thurberianum*, six replicates of *P. secunda*, and one replicate of *C. douglasii, E. nauseosa*, and *A. tridentata* to approximate a dense natural plot, and two seeds were sown into each planting position to increase the chances of seedling emergence. In cases where seedlings from both seeds established, each planting position was thinned to one seedling based on a random coin flip. To augment direct seeding, plants of each species and site were also grown in the greenhouse to serve as replacements for plants failing to establish directly from seed.

We created two treatment types, sympatric or allopatric, with six different combinations for each treatment. In the sympatric treatments, seeds of all six species from one of the six locations were sown together. In the allopatric treatments, seeds from each species and location were randomly assigned to one of six allopatric mixes, with one representative species from each location in each allopatric mix. This led to a total of 12 unique communities, which were each planted with seven or eight replicates, for a total of 90 mesocosms.

Mesocosm soil was initially watered to maximum water holding capacity and lightly watered once weekly if no natural precipitation had occurred. Each mesocosm was monitored for seedling emergence from November 2016 to May 2017, and seedlings were thinned as needed. In May 2017, for any plants that failed to recruit from seed, individuals were transplanted from seeds sown in the greenhouse. At the end of August 2017, when many of the perennial grasses had become senescent, every plant was assessed for height, crown size (length × width), senescence index (a visual estimate using a continuous integer between 0-3; 0 = no live tissue and 3 = >75% green tissue), and the number of inflorescences. Starting in late September 2017, the second growing season, a new round of monitoring began following several rainstorms. We tracked new leaf growth and survival through May 2018. As in the previous year, in May 2018, we replaced any dead individuals. From May to October 2018, we noted whether plants had green tissue (0/1) on a weekly basis and used this information to estimate the number of days a given plant had potentially photosynthetically active tissue during the growing season. At the end of August 2018, each plant was assessed for crown size, height, senescence index, and the number of inflorescences.

In October 2018, each mesocosm was invaded with 130 *B. tectorum* seeds to test invasion resistance, with seeding density based on a field survey of seed production in a moderately invaded site. Seeds were lightly raked into the soil surface, in interspaces between plants. All plants were grown through August 2019 and watered as needed as in previous years. In August 2019, the aboveground biomass of all plants was harvested, oven-dried, and weighed.

Annual productivity was estimated from measurements of native plant volume in years 1 and 2 (calculated as crown size multiplied by plant height) and from aboveground biomass in 2019, and reproductive output was estimated as the number of inflorescences per plant. Overall survival was measured from the number of transplants used to replace dead plants required at any planting location in a mesocosm over all three growing seasons. The total mass of *B. tectorum* per mesocosm was used as the indicator of invasion resistance. For species that have senescence as part of their life history strategy (all but *A. tridentata*), phenology was evaluated in two different ways: 1) the total number of days that a given plant had green tissue from fall through spring (September through May), or 2) the senescence index measured in August.

### Statistical analysis

We first used community composition models to analyze overall community composition and plant survival in each year, using linear models implemented in R version 4.1.0 (R Core Team, 2021). These models included species, community, and a species × community interaction as fixed effects with survival, plant volume, or biomass as response variables. Response variables were transformed as needed to better fit model assumptions (Table S1). *F* statistics and *p*-values were calculated using the R package ‘car’ (Fox & Weisberg, 2019). The goodness of fit (R^2^-values) was calculated with the R package ‘MuMIn’ (Barton, 2020) for this and all glm models and post-hoc contrasts were generated using the Tukey honest significant differences method with the ‘agricolae’ R package (de Mendiburu & Yaseen, 2020). Residual histograms and scatterplots of predictors and response variables were created to verify model assumptions of normality (e.g., Figure S3), homogeneity of variances (e.g., Figure 2), and linearity of relationships (e.g., Figure 4).

**Figure 1.**
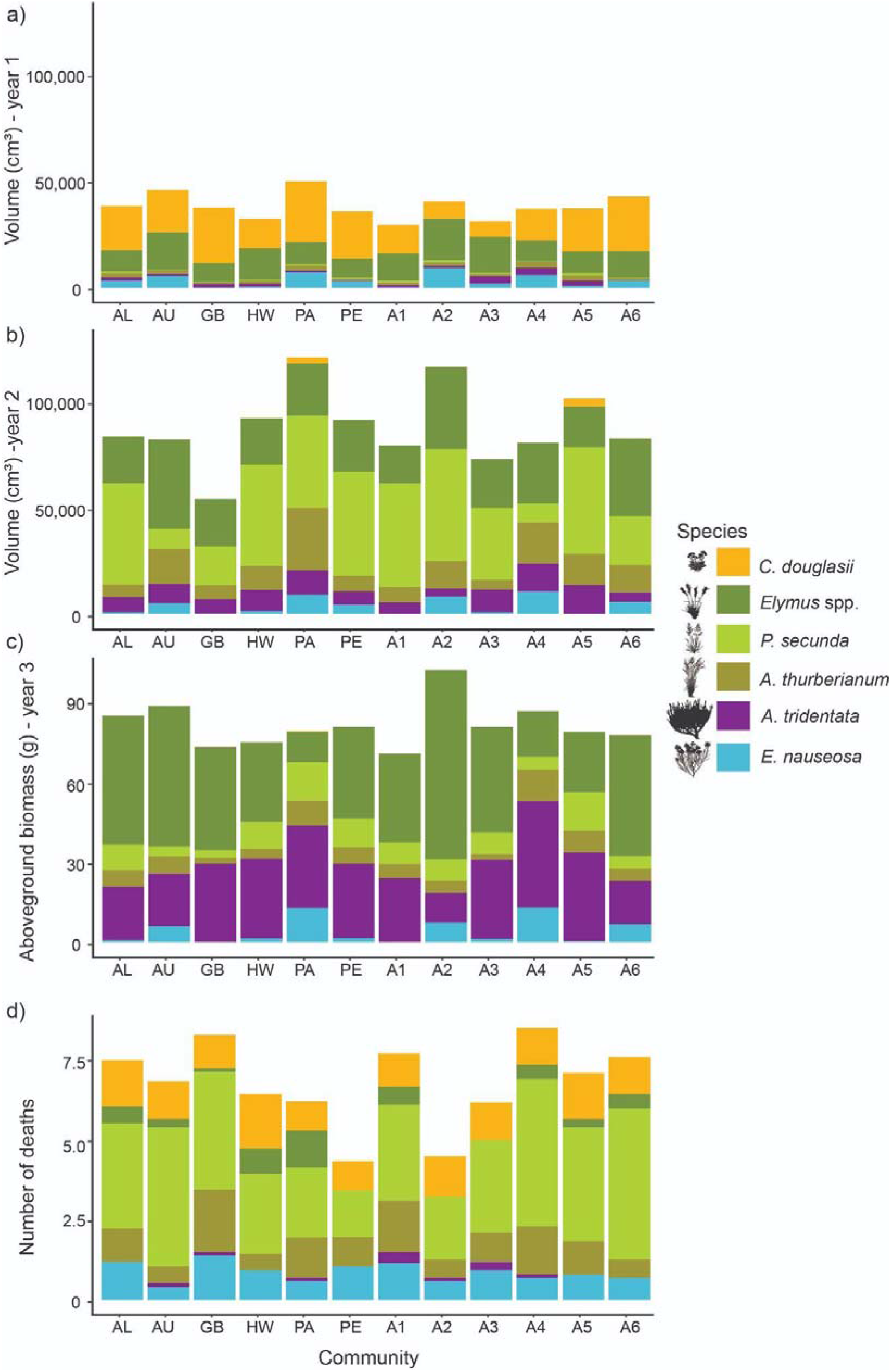
Community composition in experimental mesocosms for three growing seasons (a-c) and (d) the distribution of native plant deaths across species and communities. Volume (length × width × height) and biomass values are the averages across 7-8 mesocosms of each community type, shown without variation for visual clarity. The number of deaths shown is proportional to the number planted for each species, calculated as the total number of deaths divided by the number planted per species per mesocosm. The composition of allopatric mixes is shown in Figure 3d and abbreviations indicate the seed source (Figure S1): AL(Alturas), AU (Austin), GB (Grey’s Butte), HW (Highway 140), PA (Patagonia), PE (Peavine), A (allopatric mixtures 1-6).

**Figure 2.**
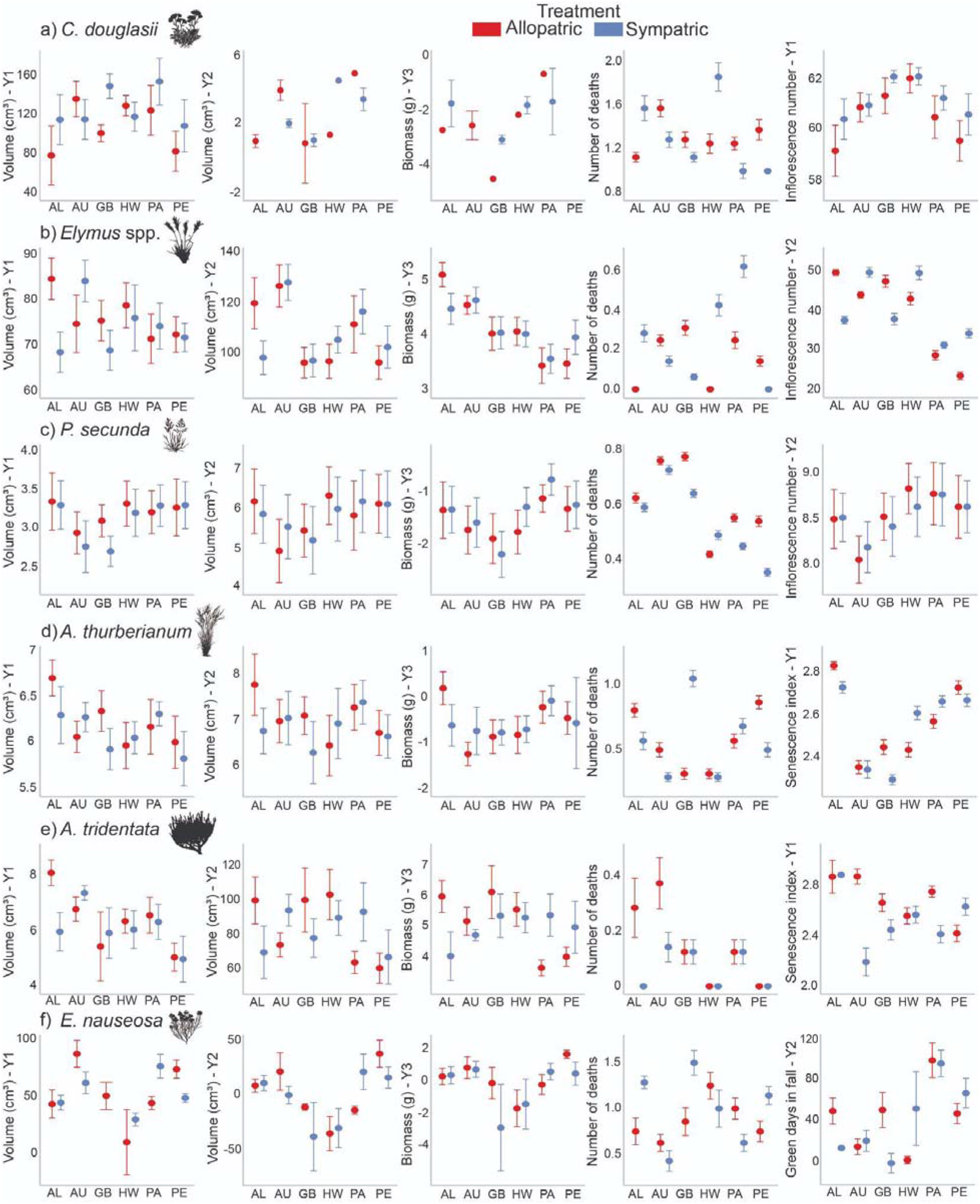
Differences in responses between sympatric and allopatric treatments for all species (a-f) and collection sites (x axis) with volume, biomass, survival across three growing seasons, and flower number or phenology. Some populations of some species had different responses to sympatric vs. allopatric neighbors, while many did not. Values are means ± SE from the species-specific models that account for the effects of plant age and mesocosm (for grasses), which were transformed as needed for analysis and are shown here without transformation (Table S1, Figure S3).

**Figure 3.**
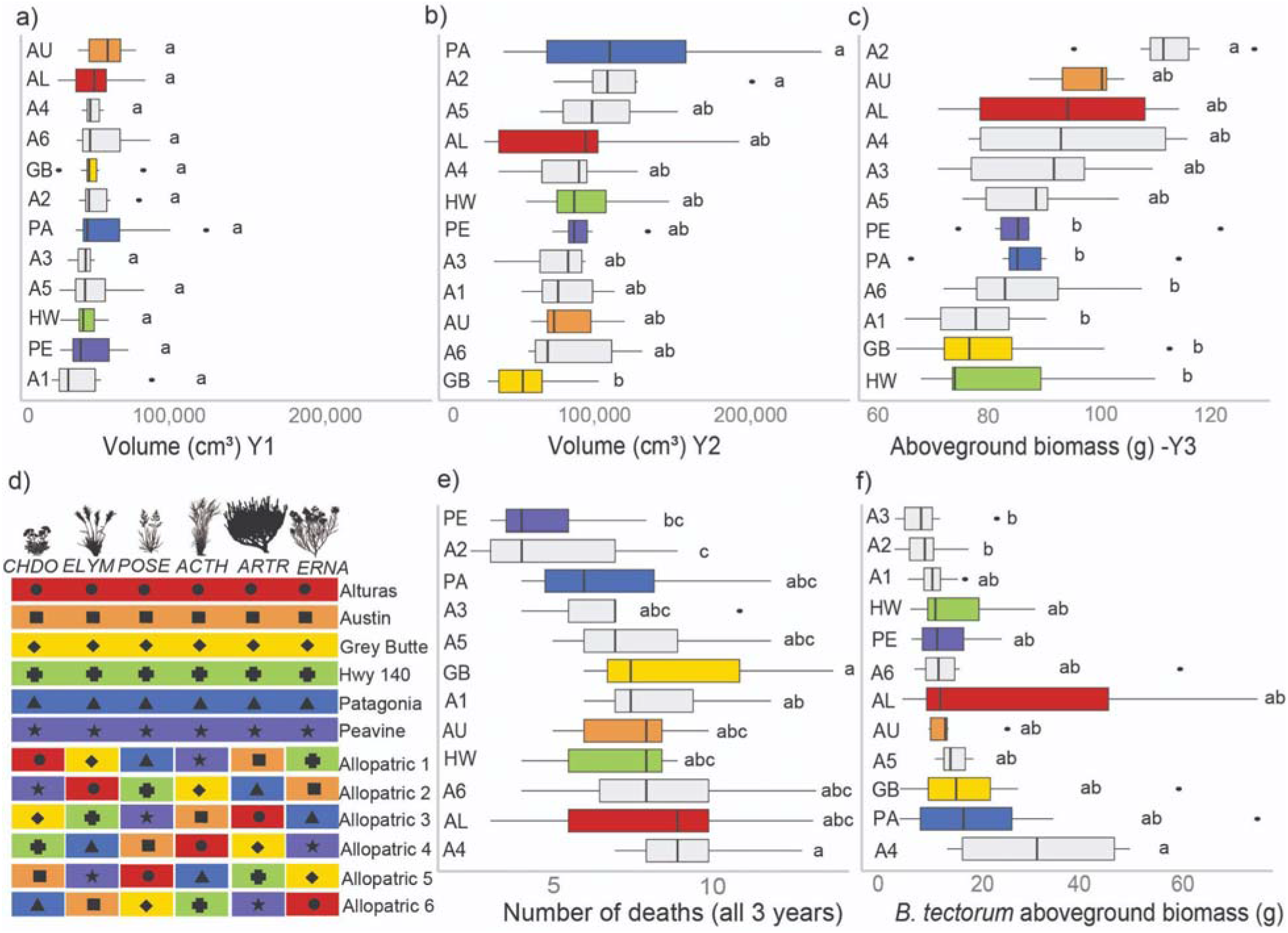
Differences among 12 unique communities in size, survival, and invasion resistance.Native plant size is represented by total plant volume (length x width x height) in the (a) first and (b) second growing season, and by (c) total aboveground biomass in the third season. The unique community design (d) is displayed along with (e) total mortality of all native plants in each mesocosm and (f) *B. tectorum* aboveground biomass from invaded mesocosms. Box plots indicate medians, first to the third quartiles, and outliers shown as black points, calculated from 7-8 replicate mesocosms per unique community; values were transformed as needed for analysis (Table S1, Figure S3).

**Figure 4.**
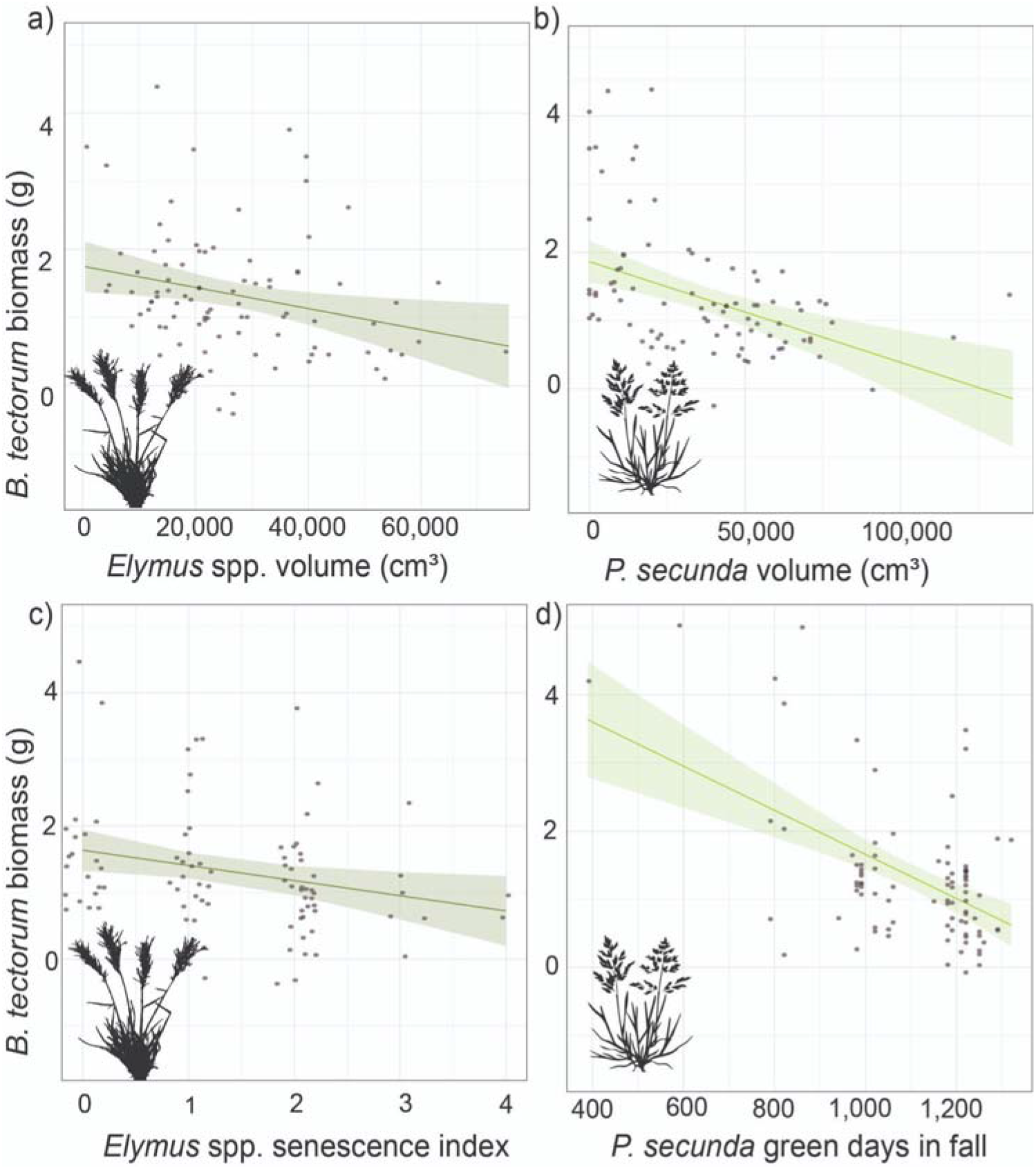
Marginal effects with pointwise 95% confidence intervals of the effects of (a) *Elymus* spp. volume, (b) *P. secunda* volume, (c) *Elymus* spp. senescence index (measurement of greenness in August), and (d) *P. secunda* phenology (the number of live green days in the fall preceding invasion) on *B. tectorum* aboveground biomass, which were the four best predictors used in the best multiple regression model (R^2^=0.44); model residuals are plotted in Figure S3.

#### Q1: Do plants perform differently when grown with sympatric or allopatric neighbors, and do responses differ among species or populations?

We created a main effects model asking whether individual plants had responses to sympatric or allopatric neighbors, using linear models with species, collection site, and treatment (allopatric or sympatric) as fixed effects. For grass species, we used linear mixed models that included mesocosm as a random (intercept) effect, as they had multiple replicates in a mesocosm (Bates et al., 2015). Response variables included survival, size, reproduction, and phenological variables. Based on our previous work, we expected considerable variation among species, thus we also built species-specific models asking whether there were species- or location-specific differences in response to sympatric or allopatric neighbors. First, we first created linear mixed models for each species that included survival as response and collection site, treatment, and the interaction between site and treatment as fixed effects. Then, we asked whether surviving plants of each species differed in size, phenology, or reproduction in allopatric or sympatric communities using the same model structure described above, but also included plant age (days since transplanting) as a covariate; response variables were transformed as needed (see Table S1 for transformations and Figure S3 for histograms of residual error).

#### Q2: Do community-level responses differ among allopatric or sympatric treatments?

We then quantified whether community-level responses (total productivity, total survival, inflorescence production, or *B. tectorum* suppression) differed among unique communities or by allopatric/sympatric treatment using linear models on the aggregated community level values per mesocosm. Total responses were calculated by summing response values across all individual plants in each mesocosm. Models included treatment (allopatric/sympatric) and unique community (one of the 12 allopatric or sympatric combinations, nested within treatment) as fixed effects, and separate models were created for each response variable and each year of measurement, with transformations as needed (Table S1, Figure S3). Means were again compared using the Tukey honest significant differences method for multiple comparisons.

#### Q3: Can we identify factors that can predict differences in invasion resistance?

After assessing whether communities differed in key performance metrics, we sought to identify factors that might explain the observed differences in *B. tectorum* invasion resistance. For this, we asked whether any individual species was having an outsized predictive effect on *B. tectorum* biomass. We took a tiered model building approach. First, we considered potential explanatory variables (survival, inflorescence production, volume, senescence level of native plants prior to invasion, and the number of live green days experienced during the fall-winter of invasion) separately for each species. For grass species, we summed the values across the replicates within a community to represent the total performance of a species. We ran a set of generalized linear models separately for each metric of performance and each species, considering whether any individual response for an individual species predicted *B. tectorum* biomass, log transformed for analysis (Table S1, Figure S3). We then retained a subset of variables for a multiple regression model, which included the most impactful explanatory variables (*p* < 0.05) from the previous analyses, further reducing as needed to ensure no variable in the model had a VIF greater than 3 (Fox & Weisberg, 2019). Finally, we retained significant variables to create a final multiple regression model of species-level responses that best explained *B. tectorum* biomass.

## Results

Community composition changed dramatically over time, consistent with successional patterns in these plant communities (Figure 1). In the first growing season, the community composition model revealed differences in the size of each species (*F*_5,468_ = 73.3, *p* <0.001), with *C. douglasii* and/or *Elymus* spp. being the largest components of experimental communities. There was also a significant community × species effect (*F*_55,468_ = 1.5, *p* = 0.015), primarily caused by several sources of *C. douglasii* growing significantly larger or smaller than collections from other locations (Figure 1a). In the second growing season, there were again differences among species (*F*_5,468_ = 76.8, *p* <0.001); *C. douglasii* almost completely died and *P. secunda* started to dominate alongside *Elymus* spp. Communities also began to differ significantly in overall volume (Figure 1b; *F*_11,468_ = 2.0, *p* = 0.030), and species behaved differently across community mixtures (community × species effect; *F*_55,468_ = 2.5, *p* <0.001), driven mostly by a few populations of *P. secunda* varying greatly among communities. In the final year, species again differed in size (Figure 1c; *F*_5,468_ = 163.6, *p* <0.0001), across community mixtures (community x species effect; *F*_55,468_ = 4.9, *p* <0.001), and in most communities, *Elymus* spp. and *A. tridentata* dominated the aboveground biomass.

In addition to differences in size, species also differed in overall survival in the community composition model, with *Elymus* spp. and *A. tridentata* experiencing the lowest mortality (Figure 1d; *F*_5,468_ = 118.9, *p* <0.001). Communities differed in overall survival (*F*_11,468_ = 2.5, *p* =0.005), with plants growing in the single-origin, sympatric Peavine communities and random, allopatric community mixture 2 having significantly greater survival overall. Finally, there were also species x community interactions, driven primarily by *P. secunda* from Austin and Grey’s Butte having a large number of deaths (*F*_55,468_ = 2.2, *p* <0.001), relative to *P. secunda* collected from other locations.

### Q1: Do plants perform differently when grown with sympatric or allopatric neighbors, and do responses differ among species or populations?

In the main effects model, species strongly differed in every response variable, and collection sites varied in all but 4 of 10 responses, but no response variable had an overall main effect of allopatric or sympatric treatments (Table 1).

**Table 1.**
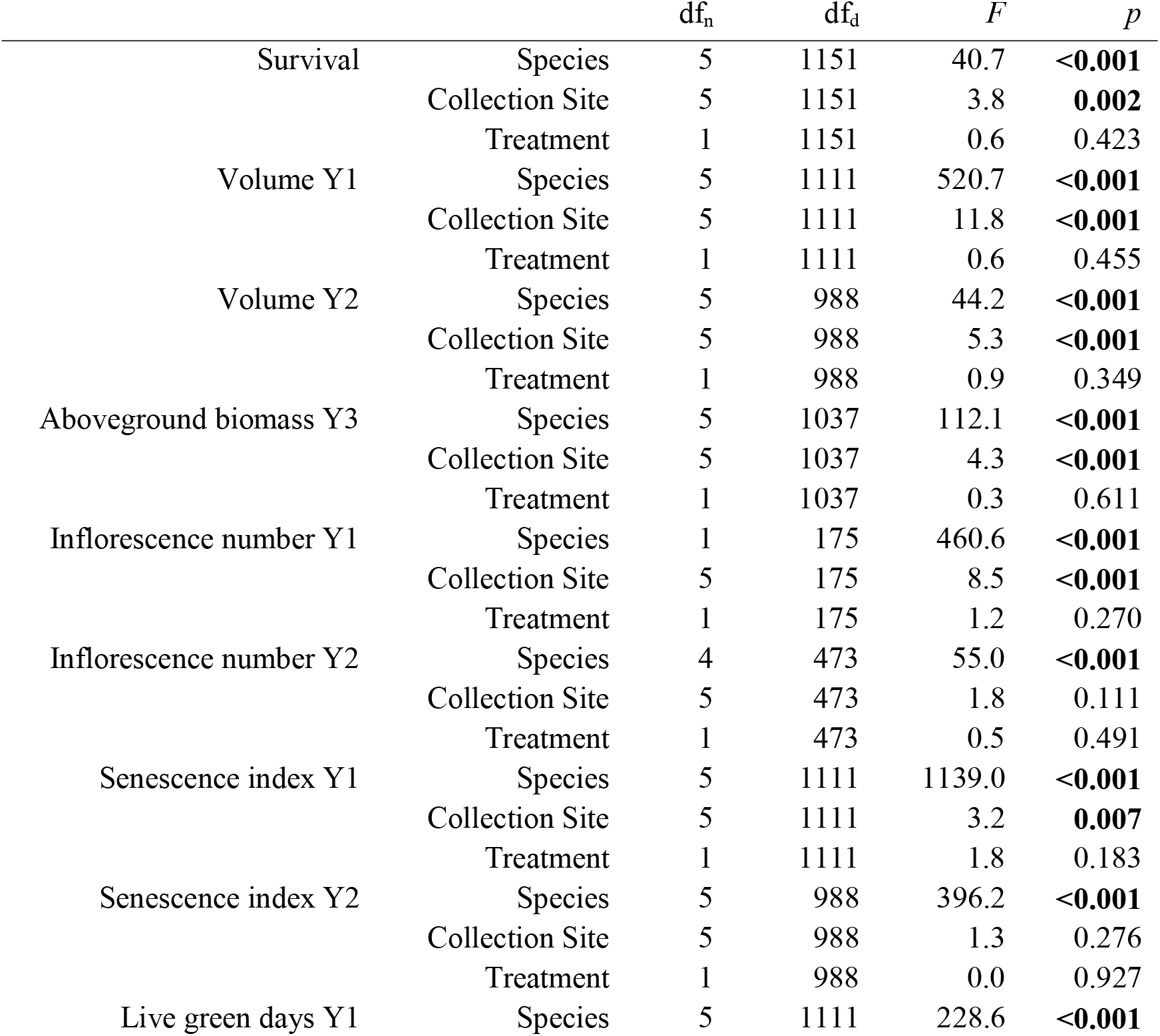
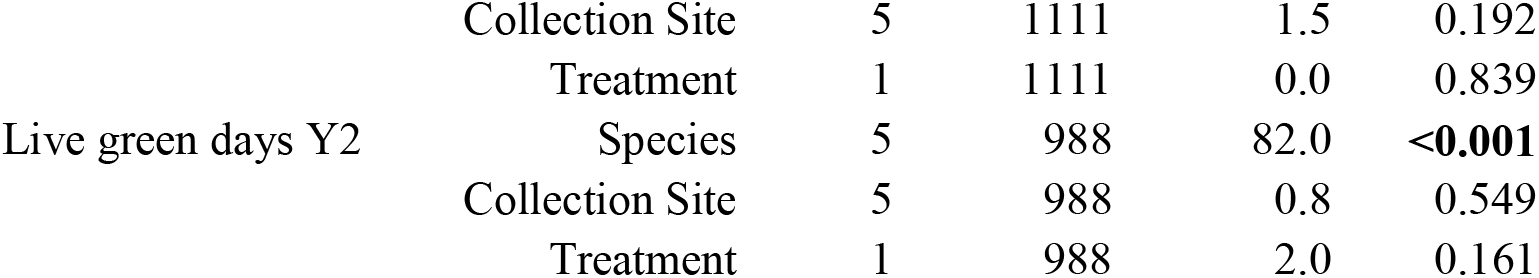
Results of main effects models showing differences among species, collection sites, and allopatric or sympatric treatments, considering survival for each planting location across the three-year experimental period, and the size (length × width × height volume measurements in years 1 and 2, and aboveground biomass in year 3), reproduction (inflorescence number), and two measures of growing season phenology of surviving plants separately for each year. Values reported are from linear models, and include numerator (n) and denominator (d) degrees of freedom (df), test statistics (*F*), and significance (*p*) with bolded values indicating significance <0.05.

Results were similar when we considered each species on its own in the species-specific model: the identity of neighboring plants within the community (allopatric or sympatric) affected response variables only for certain species from certain collection sites (Figure 2, Table S2). For example, *Elymus* spp. from Alturas benefitted from allopatry in all three growing seasons, growing 50% larger on average in the allopatric mix in the first growing season than in the single-origin, sympatric mesocosms (Figure 2b). *C. douglasii* was the only plant to experience an overall treatment effect: individuals growing in sympatric communities produced significantly more inflorescences than when grown in allopatric communities, though effects for any single collection site were generally small (Figure 2a, Table S2B).

Many response variables varied across collection locations, as either main location effects or interactions with treatment, with a few exceptions for productivity (i.e., year 2 volume in *A. thurberianum, A. tridentata*, and *C. douglasii*) and indicators of phenology (i.e., senescence index and live green days in the fall; Table S2). Only *P. secunda* varied significantly in survival based on collection location and *Elymus* spp. had a significant treatment by collection location effect, with plants sourced from Patagonia and Highway 140 experiencing greater mortality in sympatric mesocosms (Table S2A). Plant age was significantly predictive of most response variables, except for some of the phenology measurements for grass species (Table S2).

### Q2: Do community-level responses differ among allopatric or sympatric treatments?

The main effect of sympatric or allopatric treatment had no significant predictive effect for nearly all response variables in the community response model (Table 2); the exception was the number of inflorescences in year one, which was driven by the weak but overall positive flowering response of *C. douglasii* to sympatric neighbors discussed above. Unique communities varied in total productivity in the final two growing seasons, as well as total survival and *B. tectorum* biomass (Figure 3, Table 2). For instance, allopatric mixture 2, which contained the *Elymus* spp. from Alturas that had a strong positive response to sympatric neighbors (Figure 1c, Figure 2b), grew the largest native plants in the final growing season and had some of the lowest overall mortality and smallest *B. tectorum*. However, other allopatric community combinations (e.g., allopatric mixtures 4 and 6) ranged from low to average in several performance metrics (Figure 3). Similarly, sympatric communities were found in both the highest and lowest categories for each performance metric (Figure 3).

**Table 2.** Community response models showing differences in community-level survival, native plant size (volume or aboveground biomass), invader size (*B. tectorum* aboveground biomass) among twelve unique communities and two treatments (allopatric or sympatric). Values reported are from linear models, and include numerator degrees of freedom (df), test statistics (*F*), and significance (*p*) with bolded values indicating significance <0.05. The denominator degrees of freedom are 78 across all models.

|  |  | df<sub>n</sub> | *F* | *p* |
| --- | --- | --- | --- | --- |
| Survival | Community | 10 | 3.0 | **0.003** |
|  | Treatment | 1 | 0.5 | 0.502 |
| Volume Y1 | Community | 10 | 1.2 | 0.287 |
|  | Treatment | 1 | 0.7 | 0.405 |
| Volume Y2 | Community | 10 | 2.2 | **0.026** |
|  | Treatment | 1 | 0.9 | 0.345 |
| Native biomass Y3 | Community | 10 | 3.6 | **<0.001** |
|  | Treatment | 1 | 1.1 | 0.289 |
| *B. tectorum* biomass | Community | 10 | 2.2 | **0.024** |
|  | Treatment | 1 | 1.2 | 0.286 |
| Inflorescence no. Y1 | Community | 10 | 2.8 | **0.006** |
|  | Treatment | 1 | 4.1 | **0.047** |
| Inflorescence no. Y2 | Community | 10 | 4.2 | **<0.001** |
|  | Treatment | 1 | 0.1 | 0.712 |

While both biomass and invasion suppression differed among communities (Figure 3, Table 2), the variation in native biomass production among communities was much lower than the variation in *B. tectorum* suppression. For example, in the 3^rd^ year, there was a ~2-fold difference between the smallest and largest native biomass (60 g vs. 128 g), while there was an ~83-fold difference between the smallest and largest *B. tectorum* biomass (0.07 g vs 5.78 g), and a large range in the number of *B. tectorum* that survived in each mesocosm (3-51, average ~28 plants).

In year two when all three grass species flowered, flower number varied by community, with allopatric mix 2 having the most flowers and allopatric mixture 4 having the least (Table 2, results not shown). Phenology and greenness timing were the responses least affected by either treatment or community, with only senescence index varying significantly among communities in the first year, driven by allopatric mixture 2 having significantly less senescence than allopatric mix 3 (results not shown).

### Q3: Can we identify factors that can predict differences in invasion resistance?

We considered the relative importance of multiple factors that may contribute to *B. tectorum* suppression. Models including individual response variables for each species had variable predictive abilities (range of R^2^ = 0 - 0.20) and revealed that three species influenced *B. tectorum* invasion: *Elymus* spp., *P. secunda*, and, less so, *A. tridentata* (Table S3). Through our tiered model building approach, we found the most predictive model (R^2^ =0.44) included the volume of *Elymus* spp. and *P. secunda*, the *Elymus* spp. senescence index (a measure of live green tissue present in August), and *P. secunda* green days over the fall-spring during invasion (Figure 4).

Greater *Elymus* spp. volume (*F*_1,85_ =10.4,*p* =0.002) and *P. secunda* volume (*F*_1,85_ =19.3, *p* <0.001) were associated with the smallest *B. tectorum* (Figure 4a,b), and *Elymus* spp. with higher senescence indices (more green tissue) in August were significantly correlated with lower *B. tectorum* biomass (*F*_1,85_ =8.9, *p* =0.004; Figure 4c). Mesocosms containing *P. secunda* with fewer live green days over the fall had larger *B. tectorum* (*F*_1,85_ =14.0, *p* <0.001; Figure 4d).

## Discussion

Interactions among plants are fundamental components of all terrestrial ecosystems, and yet our understanding of how evolutionary and geographic history affect the nature of plant-plant interactions and community function is still incomplete. While positive effects of sympatric history have been observed with small numbers of interacting species (Aubree et al., 2020; Chanway et al., 1988; Grady et al., 2017; van Moorsel et al., 2018), our study provides the most species-rich test to-date of the effects of sympatry on community functions from wild, unmanipulated plant communities. Our results demonstrate that while community composition can sometimes affect plant performance and emergent community properties, planting sympatric communities is not sufficient to ensure high ecosystem services (Table 1). Where others have found clear benefits of growing plants with sympatric neighbors (Camarretta et al., 2020; Grady et al., 2017; van Moorsel et al., 2018), we instead found more nuanced results that varied across species, over time, and among response variables (Figure 2). Though responses were idiosyncratic, it is notable that one of the communities with the greatest resistance to invasion was formed when a population of the highly influential species (*Elymus* spp.) responded favorably to allopatric neighbors (*Elymus* spp. from Alturas; Figures 1 and 2). Thus, while our results do not support the idea that a history of sympatry produces complementarity in resource use that leads to improved individual plant fitness and reduced opportunity for species invasion, our results do confirm that mixtures of species from different sources can result in altered community properties.

The lack of consistent response to sympatric or allopatric neighbors observed here supports the idea that the processes of niche differentiation or ecological sorting can be context dependent (Eisen & Geber, 2018). Sympatric communities did not display more complete use of resources, as evidenced by no overall aboveground biomass advantage, greater resistance to *B. tectorum*, greater survival, or greater reproduction for most species (Figure 3). We do note that the annual/biennial forb, *C. douglasii* did have a positive overall response to sympatric neighbors in an important fitness metric, inflorescence number, but it was the only consistent response. It is possible that productivity is not always enhanced through co-adaptation and instead of actively suppressing invasion, plants may adapt to tolerate invasive species, resulting in more stable, albeit less productive, communities (Aubree et al., 2020). We did not measure tolerance to invasion in this experiment, but it is possible that quantifying declines in native plant growth or reproduction in response to invasion could reveal differences in allopatric/sympatric communities that were not demonstrated by measuring productivity or invasive suppression alone. We also note our experimental communities consisted of mid- to long-lived species, and while our three-year experiment was longer than some, it is likely that additional dynamics would be revealed in a study of older plants. Indeed, differences in productivity among communities became more pronounced over the three-year study period (Figure 3).

Interactions among multiple species within each mesocosm can also create complex coexistence networks where higher-order interactions occur (Landi et al., 2018; Levine et al., 2017). In our experiment, it may be that the density and spatial arrangement of plants within mesocosms released some species from intraspecific competition, and instead emphasized interspecific interactions. This may account for some of our findings if a particularly dominant species or population benefited through release from competition with intraspecific neighbors (Martínez-Blancas & Martorell, 2020). Additionally, it is possible that these plants are not interacting in the wild as much as one might assume, and that their sympatric co-occurrence is not associated with reciprocal selection pressures (Hubbell, 2001). Because we sampled wild communities, we do not know the length of interaction history, which is a factor that others have deliberately incorporated into manipulative experiments assessing the impacts of evolutionary history on community assembly (Weisser et al., 2017). Further, we note that while our experiment contained a relatively large number of species relative to pair-wise experiments, we only sampled a few community dominants, and acknowledge that natural plant communities would contain many more species of variable size, distribution, and age, all of which could affect community characteristics not captured here.

While the strength of interactions between the wild plants utilized in our experiment is unknown, there is clear evidence of interactions between native plants and invasive species in the Intermountain West (Colautti & Lau, 2015; Leger, 2008; Leger & Espeland, 2010) and clear evidence of interactions between native plants and invasives in our experiment (Table S3). For example, we found that the size and phenology of two grass species (*Elymus* spp. and *P. secunda*) more strongly predicted *B. tectorum* suppression than the community’s overall native plant volume and senescence (Figure 4). The mesocosms with the lowest *B. tectorum* biomass were dominated by these two native grass species, and we note that mesocosms that had more even biomass distribution across the six species had low (allopatric mix 4) to average (Patagonia sympatric community) *B. tectorum* suppression, indicating that while they represented a more even native community, they were not as effective at resisting invasion. In restoration contexts, this leads to the question of how to best create a community when we value opposing ecosystem properties, such as biodiverse habitats versus simpler communities with greater invasion resistance (e.g., Davies & Johnson 2017). It is possible that balancing these values will require careful consideration of not only species identities but species origins as well.

Few of our results aligned with expectations based on previous studies in grasslands (van Moorsel et al., 2018; Zuppinger-Dingley et al., 2014) and work within the same functional groups (Camarretta et al., 2020; Grady et al., 2017). It is possible that within our semi-arid study system, abiotic factors are more important drivers of the evolution of plant form, and the coevolutionary dynamics observed in more mesic systems are secondary influences within these plant communities. When considering the important community outcome of invasion suppression, we found evidence to support the ‘dominance’ effect theory since certain populations of *Elymus* spp. and *P. secunda* had strong effects on *B. tectorum* biomass (Figure 4; Mahaut et al., 2020). Further, while the sympatric/allopatric origin of plant neighbors had very few effects on individual plant performance, we were able to assemble one allopatric plant community where unknown factors aligned to increase the growth of one important native species and provided outsized suppression effects. Despite the fact that the low productivity of our system likely limited the importance of interactions and coevolutionary history, our results are consistent with a bigger picture of complex diversity effects that has emerged in recent years. In particular, a large analysis of grassland experiments from Europe found that many of the important effects of individual species were only realized under specific combinations of conditions (e.g., certain weather in particular years) (Isbell et al., 2011). In an applied context, our results suggest that there is unlikely to be any benefit from simply planting sympatric communities when assembling new communities via ecological restoration in this system.

Instead, it may be possible to create mixtures of the same species from different source populations that can result in different ecosystem functions.

## Acknowledgments

This project was funded by grant 2017-67019-26336 from the USDA National Institute of Food and Agriculture Sustainable Agroecosystems program and Cooperative Agreement No. L16AC00318 from the Bureau of Land Management. Thank you to Drs. Kevin Shoemaker, Elizabeth Pringle, and Jenny Ouyang for discussions about design, analysis, and feedback to initial manuscript drafts. Additional thanks to Dr. Robert Nowak and Scot Ferguson for providing the mesocosms, and Shannon Swim, Owen Baughman, Meagan O’Farrell, Chase Estes, Georgia Peterson, Kalin Ingstad, Sage Ellis, Trevor Carter, Amber Durfee, Katelyn Josifko, Carley Crosby, and Casey Iwamoto for assistance with seed collection and processing, mesocosm filling and planting, and data collection.

