## Supplemental material for "Does a history of co-occurrence predict plant performance, community productivity, or invasion resistance?"

### Appendix 1: Supplemental Tables and Figures

###### Table S1. Summary of variable transformations for the main effects and species-specific models, selected to better meet model assumptions. Distribution of residuals from representative community and individual species models are shown in Figure S3. “-” indicates that this response was not analyzed at the community level or for a given species, either because all values were zero, values were invariant, or values were highly non-normally distributed despite transformation. Species abbreviations are as follows: *C. douglasii* (CHDO), *Elymus* spp. (ELYMU), *P. secunda* (POSE), *A. thurberianum* (ACTH), *A. tridentata* (ARTR), and *E. nauseosa* (ERNA).

|  | Community | CHDO | ELYMU | POSE | ACTH | ARTR | ERNA |
| --- | --- | --- | --- | --- | --- | --- | --- |
| Survival | Square root | None | None | None | None | None | None |
| Volume Y1 | Log | Square root | Square root | Log | Log | Log | Square root |
| Volume Y2 | Log | Log | Square root | Log | Log | Square root | Square root |
| Biomass Y3 | Log | Log | Square root | Log | Log | Square root | Log |
| Inflorescence no. Y1 | Square root | Log | Log | - | - | - | - |
| Inflorescence no. Y2 | Square root | - | None | Log | Log | - | - |
| Direct Emergence | - | None | None | None | None | - | None |
| Senescence index Y1 | Square root | None | None | - | None | None | None |
| Senescence index Y2 | None | None | None | - | None | None | None |
| Live green days Y1 | None | - | - | - | - | - | None |
| Live green days Y2 | Square root | - | None | - | None | - | None |
| *B. tectorum* biomass | Log |  |  |  |  |  |  |

###### Table S2. Results of species-specific linear models reveal differences among allopatric and sympatric treatments (Treat.), communities (Comm.), and the treatment x community interaction (T x C) in analyses conducted separately for each species of grasses (A) and forb/shrubs (B). Age is included as a continuous variable for all responses except survival. Values reported for the grass species are from linear mixed models that included mesocosm as a random factor since multiple individuals were planted for each species in a mesocosm; these results include degrees of freedom (df) and chi-squared test statistics, and significance (*p*) with bolded values indicating significance <0.05. Values reported for the shrubs and *C. douglasii* are from linear models and include numerator degrees of freedom, test statistics (*F*) in addition to *p* indicating significance*.* “-” indicates variables either not measured, lacking variability (e.g., *P. secunda* did not flower in its first growing season and all *A. tridentata* remained green throughout the fall), or not analyzed due to non-normal residuals and unequal variance despite transformation (specifically, the live green days response was not analyzed for any grass taxa in year 1 or for *P. secunda* in year 2 due to highly non-normal distributions).

| A) |  | *Elymus* spp. | | | | *P. secunda* | | | | *A. thurberianum* | | | |
| --- | --- | --- | --- | --- | --- | --- | --- | --- | --- | --- | --- | --- | --- |
|  |  | df | | ꭓ^2^ | *p* | df | | ꭓ^2^ | *p* | df | | ꭓ^2^ | *p* |
| Survival | Treat. | 1 | | 1.6 | 0.206 | 1 | | 2.4 | 0.123 | 1 | | 0.0 | 0.913 |
|  | Comm. | 5 | | 9.1 | 0.105 | 5 | | 27.0 | **<0.001** | 5 | | 7.7 | 0.173 |
|  | T x C | 5 | | 12.8 | **0.026** | 5 | | 3.2 | 0.670 | 5 | | 10.7 | 0.059 |
| Volume Y1 | Treat. | 1 | | 1.2 | 0.283 | 1 | | 3.8 | 0.052 | 1 | | 2.4 | 0.121 |
|  | Comm. | 5 | | 6.9 | 0.228 | 5 | | 62.3 | **<0.001** | 5 | | 32.4 | **<0.001** |
|  | T x C | 5 | | 13.6 | **0.018** | 5 | | 10.0 | 0.075 | 5 | | 14.0 | **0.015** |
|  | Plant age | 1 | | 16.5 | **<0.001** | 1 | | 240.6 | **<0.001** | 1 | | 72.9 | **<0.001** |
| Volume Y2 | Treat. | 1 | | 0.0 | 0.975 | 1 | | 0.0 | 0.958 | 1 | | 0.7 | 0.395 |
|  | Comm. | 5 | | 28.2 | **<0.001** | 5 | | 12.5 | **0.028** | 5 | | 6.4 | 0.268 |
|  | T x C | 5 | | 6.7 | 0.242 | 5 | | 2.4 | 0.789 | 5 | | 5.4 | 0.366 |
|  | Plant age | 1 | | 98.0 | **<0.001** | 1 | | 162.3 | **<0.001** | 1 | | 134.5 | **<0.001** |
| Biomass Y3 | Treat. | 1 | | 0.0 | 0.908 | 1 | | 0.7 | 0.395 | 1 | | 0.0 | 0.984 |
|  | Comm. | 5 | | 75.2 | **<0.001** | 5 | | 31.9 | **<0.001** | 5 | | 17.5 | **0.004** |
|  | T x C | 5 | | 9.9 | 0.077 | 5 | | 7.1 | 0.213 | 5 | | 4.8 | 0.436 |
|  | Plant age | 1 | | 119.3 | **<0.001** | 1 | | 336.4 | **<0.001** | 1 | | 154.7 | **<0.001** |
| Inflorescence no. Y1 | Treat. | 1 | | 0.0 | 0.944 | - | | - | - | - | | - | - |
|  | Comm. | 5 | | 40.2 | **<0.001** | - | | - | - | - | | - | - |
|  | T x C | 5 | | 8.8 | 0.116 | - | | - | - | - | | - | - |
|  | Plant age | 1 | | 52.1 | **<0.001** | - | | - | - | - | | - | - |
| Inflorescence no. Y2 | Treat. | 1 | | 0.0 | 0.986 | 1 | | 0.1 | 0.725 | 1 | | 0.0 | 0.864 |
|  | Comm. | 5 | | 43.1 | **<0.001** | 5 | | 28.1 | **<0.001** | 5 | | 4.4 | 0.489 |
|  | T x C | 5 | | 13.1 | 0.022 | 5 | | 1.5 | 0.907 | 5 | | 2.1 | 0.840 |
|  | Plant age | 1 | | 42.3 | **<0.001** | 1 | | 125.6 | **<0.001** | 1 | | 47.1 | **<0.001** |
| Senescence index Y1 | Treat. | 1 | | 0.2 | 0.676 | - | | - | - | 1 | | 0.1 | 0.770 |
|  | Comm. | 5 | | 23.0 | **<0.001** | - | | - | - | 5 | | 23.6 | **<0.001** |
|  | T x C | 5 | | 7.7 | 0.174 | - | | - | - | 5 | | 2.7 | 0.750 |
|  | Plant age | 1 | | 1.1 | 0.288 | - | | - | - | 1 | | 35.3 | **<0.001** |
| Senescence index Y2 | Treat. | 1 | | 0.2 | 0.679 | - | | - | - | 1 | | 0.1 | 0.814 |
|  | Comm. | 5 | | 43.7 | **<0.001** | - | | - | - | 5 | | 3.1 | 0.688 |
|  | T x C | 5 | | 4.6 | 0.471 | - | | - | - | 5 | | 3.7 | 0.588 |
|  | Plant age | 1 | | 1.1 | 0.294 | - | | - | - | 1 | | 81.3 | **<0.001** |
| Live green days Y2 | Treat. | 1 | | 0.2 | 0.671 | - | | - | - | 1 | | 1.4 | 0.236 |
|  | Comm. | 5 | | 14.9 | **0.011** | - | | - | - | 5 | | 3.4 | 0.642 |
|  | T x C | 1 | | 3.4 | 0.642 | - | | - | - | 1 | | 3.7 | 0.598 |
|  | Plant age | 5 | | 39.1 | **<0.001** | - | | - | - | 5 | | 38.3 | **<0.001** |
| B) |  | *C. douglasii* | | | | *A. tridentata* | | | | *E. nauseosa* | | | |
|  |  | df_n_ | df_d_ | *F* | *p* | df_n_ | df_d_ | *F* | *p* | df_n_ | df_d_ | *F* | *p* |
| Survival | Treat. | 1 | 78 | 0.0 | 0.980 | 1 | 78 | 1.0 | 0.312 | 1 | 78 | 0.4 | 0.537 |
|  | Comm. | 5 | 78 | 1.1 | 0.356 | 5 | 78 | 1.0 | 0.431 | 5 | 78 | 1.0 | 0.449 |
|  | T x C | 5 | 78 | 2.0 | 0.090 | 5 | 78 | 0.4 | 0.817 | 5 | 78 | 0.8 | 0.538 |
| Volume Y1 | Treat. | 1 | 75 | 2.2 | 0.139 | 1 | 74 | 0.9 | 0.333 | 1 | 56 | 0.1 | 0.726 |
|  | Comm. | 5 | 75 | 1.5 | 0.216 | 5 | 74 | 4.8 | **<0.001** | 4 | 56 | 4.6 | **0.001** |
|  | T x C | 5 | 75 | 0.8 | 0.568 | 5 | 74 | 1.8 | 0.122 | 4 | 56 | 3.5 | **0.012** |
|  | Plant age | 1 | 75 | 43.9 | **<0.001** | 1 | 74 | 28.9 | **<0.001** | 1 | 56 | 24.5 | **<0.001** |
| Volume Y2 | Treat. | 1 | 11 | 0.0 | 0.997 | 1 | 77 | 0.0 | 0.884 | 1 | 60 | 0.2 | 0.682 |
|  | Comm. | 4 | 11 | 4.4 | **0.022** | 5 | 77 | 1.5 | 0.210 | 5 | 60 | 4.4 | **0.002** |
|  | T x C | 4 | 11 | 5.5 | **0.011** | 5 | 77 | 1.7 | 0.135 | 5 | 60 | 1.9 | 0.105 |
|  | Plant age | 1 | 11 | 66.6 | **<0.001** | 1 | 77 | 27.4 | **<0.001** | 1 | 60 | 59.6 | **<0.001** |
| Biomass Y3 | Treat. | 1 | 7 | 0.5 | 0.501 | 1 | 76 | 0.1 | 0.798 | 1 | 57 | 1.8 | 0.181 |
|  | Comm. | 4 | 7 | 1.4 | 0.334 | 5 | 76 | 1.4 | 0.222 | 5 | 57 | 4.3 | **0.002** |
|  | T x C | 3 | 7 | 0.5 | 0.696 | 5 | 76 | 2.3 | **0.049** | 5 | 57 | 2.5 | **0.040** |
|  | Plant age | 7 | 7 | 5.6 | **0.050** | 1 | 76 | 31.6 | **<0.001** | 1 | 57 | 64.3 | **<0.001** |
| Inflorescence no. Y1 | Treat. | 1 | 75 | 4.8 | **0.031** | - | - | - | - | - | - | - | - |
|  | Comm. | 5 | 75 | 6.3 | **<0.001** | - | - | - | - | - | - | - | - |
|  | T x C | 5 | 75 | 0.5 | 0.801 | - | - | - | - | - | - | - | - |
|  | Plant age | 1 | 75 | 47.9 | **<0.001** | - | - | - | - | - | - | - | - |
| Direct Emergence | Treat. | 1 | 58 | 8.3 | **0.006** | - | - | - | - | - | - | - | - |
|  | Comm. | 5 | 58 | 3.6 | **0.007** | - | - | - | - | - | - | - | - |
|  | T x C | 5 | 58 | 1.2 | 0.304 | - | - | - | - | - | - | - | - |
| Senescence index Y1 | Treat. | 1 | 77 | 1.5 | 0.217 | 1 | 77 | 0.0 | 0.998 | 1 | 77 | 1.4 | 0.238 |
|  | Comm. | 5 | 77 | 1.5 | 0.186 | 5 | 77 | 1.0 | 0.403 | 5 | 77 | 1.9 | 0.096 |
|  | T x C | 5 | 77 | 1.5 | 0.188 | 5 | 77 | 0.5 | 0.801 | 5 | 77 | 3.2 | **0.011** |
|  | Plant age | 1 | 77 | 4.2 | **0.043** | 1 | 77 | 86.8 | **<0.001** | 1 | 77 | 9.0 | **0.004** |
| Senescence index Y2 | Treat. | 1 | 77 | 0.1 | 0.811 | 1 | 77 | 2.9 | 0.091 | 1 | 77 | 0.6 | 0.434 |
|  | Comm. | 5 | 77 | 0.5 | 0.784 | 5 | 77 | 0.8 | 0.553 | 5 | 77 | 2.0 | 0.093 |
|  | T x C | 5 | 77 | 1.3 | 0.288 | 5 | 77 | 1.0 | 0.441 | 5 | 77 | 0.6 | 0.691 |
|  | Plant age | 1 | 77 | 10.3 | **0.002** | 1 | 77 | 12.3 | **<0.001** | 1 | 77 | 43.7 | **<0.001** |
| Live green days Y1 | Treat. | - |  | - | - | - |  | - | - | 1 | 77 | 0.4 | 0.517 |
|  | Comm. | - |  | - | - | - |  | - | - | 5 | 77 | 2.2 | 0.065 |
|  | T x C | - |  | - | - | - |  | - | - | 5 | 77 | 0.9 | 0.478 |
|  | Plant age | - |  | - | - | - |  | - | - | 1 | 77 | 14.0 | **<0.001** |
| Live green days Y2 | Treat. | - |  | - | - | - |  | - | - | 1 | 77 | 0.3 | 0.604 |
|  | Comm. | - |  | - | - | - |  | - | - | 5 | 77 | 3.6 | **0.006** |
|  | T x C | - |  | - | - | - |  | - | - | 5 | 77 | 0.9 | 0.488 |
|  | Plant age | - |  | - | - | - |  | - | - | 1 | 77 | 1.9 | 0.172 |

###### Table S3. Results for linear models testing for factors that affected aboveground *B. tectorum* biomass, considering the effects of survival, size, or phenology of native plants within each mesocosm. Models were run separately for each species, and “-” indicates either that a particular response was inapplicable to certain species (e.g., very few individuals flowered for *A. tridentata* or *E. nauseosa*) or was invariant (all *A. tridentata* individuals were equally green, all the time). Values reported include test statistics (*F*) and significance (*p*) with bolded values indicating significance <0.05. The numerator degrees of freedom are 1 and denominator degrees of freedom are 88 across all models. All variables with bolded significance values moved into a second tier of model selection. In that analysis, only *Elymus* spp. volume, *P. secunda* volume, *Elymus* spp. senescence index, and *P. secunda* number of live green days in fall had coefficients significantly different from zero, and thus moved into the final selected model (Figure 4).

|  | Survival | | Volume Y2 | | Infl. No. Y2 | | Senescence index Y2 | | Live green days Y2 | |
| --- | --- | --- | --- | --- | --- | --- | --- | --- | --- | --- |
|  | *F* | *p* | *F* | *p* | *F* | *p* | *F* | *p* | *F* | *p* |
| *C. douglasii* | 0.2 | 0.687 | 0.9 | 0.334 | 0.5 | 0.485 | 1.3 | 0.262 | 1.8 | 0.181 |
| *Elymus* spp. | 12.5 | **<0.001** | 12.0 | **<0.001** | 8.3 | **0.005** | 6.9 | **0.010** | 4.4 | **0.040** |
| *P. secunda* | 9.1 | **0.003** | 22.2 | **<0.001** | 7.6 | **0.007** | 0.0 | 0.944 | 18.3 | **<0.001** |
| *A. thurberianum* | 0.5 | 0.485 | 0.1 | 0.806 | 0.2 | 0.672 | 0.8 | 0.388 | 0.0 | 0.964 |
| *A. tridentata* | 0.0 | 0.949 | 4.0 | **0.049** | - | - | - | - | - | - |
| *E. nauseosa* | 0.0 | 0.866 | 0.6 | 0.440 | - | - | 0.1 | 0.718 | 1.5 | 0.225 |

###### Figure S1. Six collection sites (red circles) were included for all species and one planting site (blue triangle) was established.

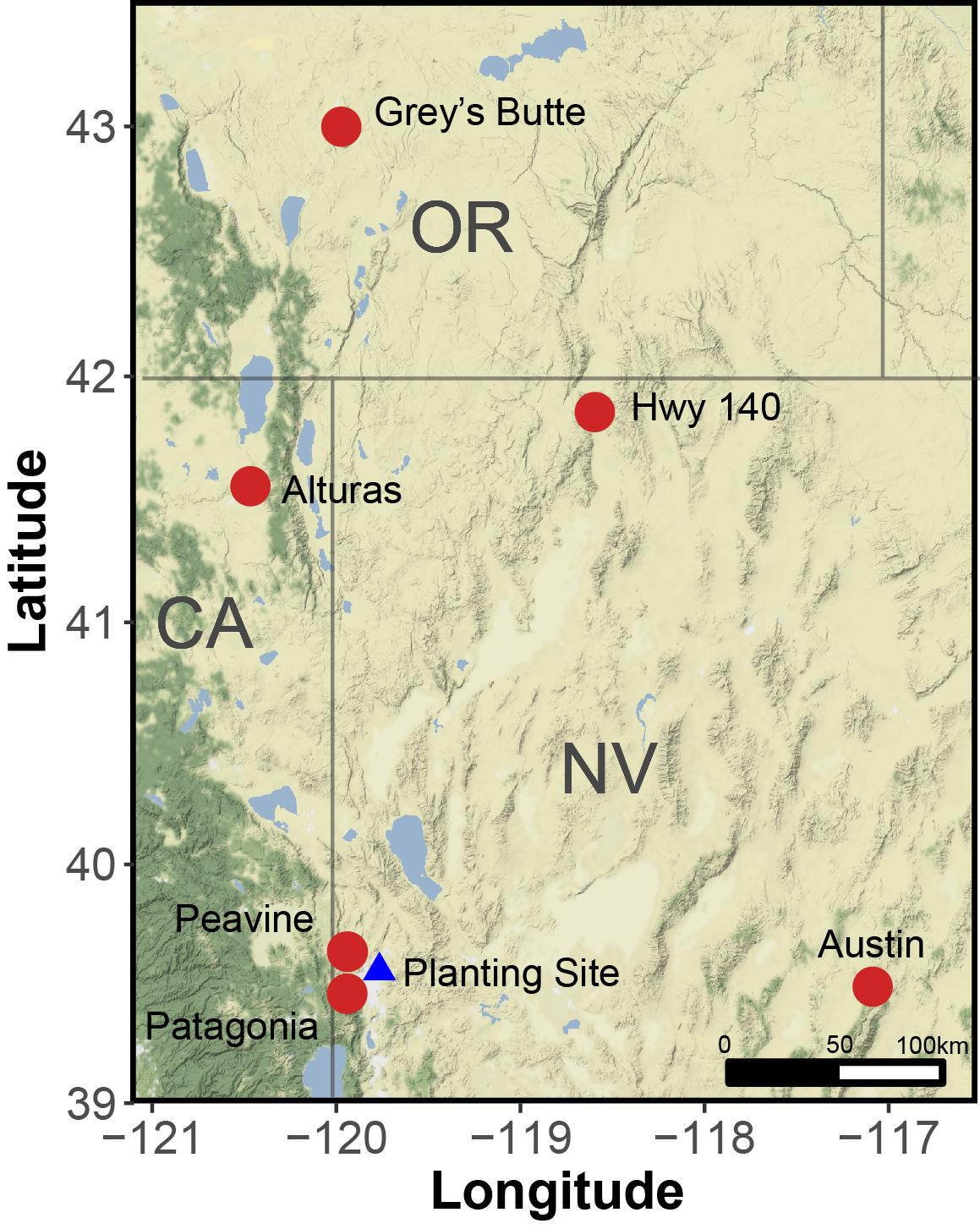

Figure S2. Experimental design showing the number and arrangement of each species in every mesocosm and an example mesocosm at four months after establishment.

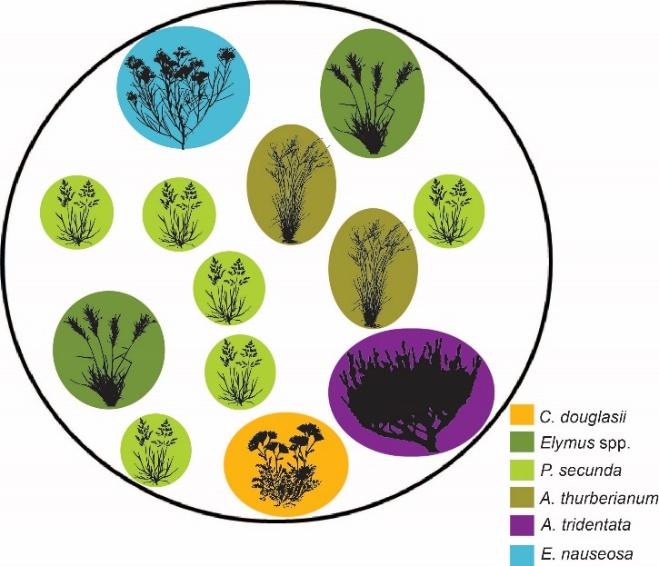

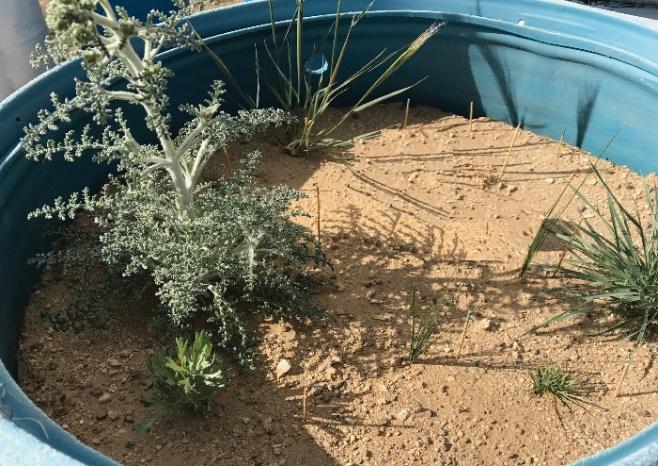

Figure S3. Distribution of model residuals for either community-level analyses (first column) or species-specific models (all other columns), after transformations as described in Table S1; the “-“ in this figure correspond to “-“ in Table S1. Residuals for species-specific models correspond to Q1, with species, collection site, treatment (allopatric or sympatric), and plant age (continuous variable) as fixed effects, corresponding to Figure 2 in the main text. Residuals for community models correspond to Q2, for models containing: treatment (allopatric or sympatric) and unique community (one of the 12 allopatric or sympatric combinations, nested within treatment) as fixed effects, corresponding to Figure 3. Finally, the distribution of residuals of *B. tectorum* biomass are shown for Q3, for the model containing the volume of *Elymus* spp. and *P. secunda*, the *Elymus* senescence index, and *P. secunda* green days over the fall-spring during invasion, corresponding to Figure 4. Residuals for other community and species-specific models reported in text and tables were very similar, as they used the same transformations and only slightly different model structures, and are thus not shown here.

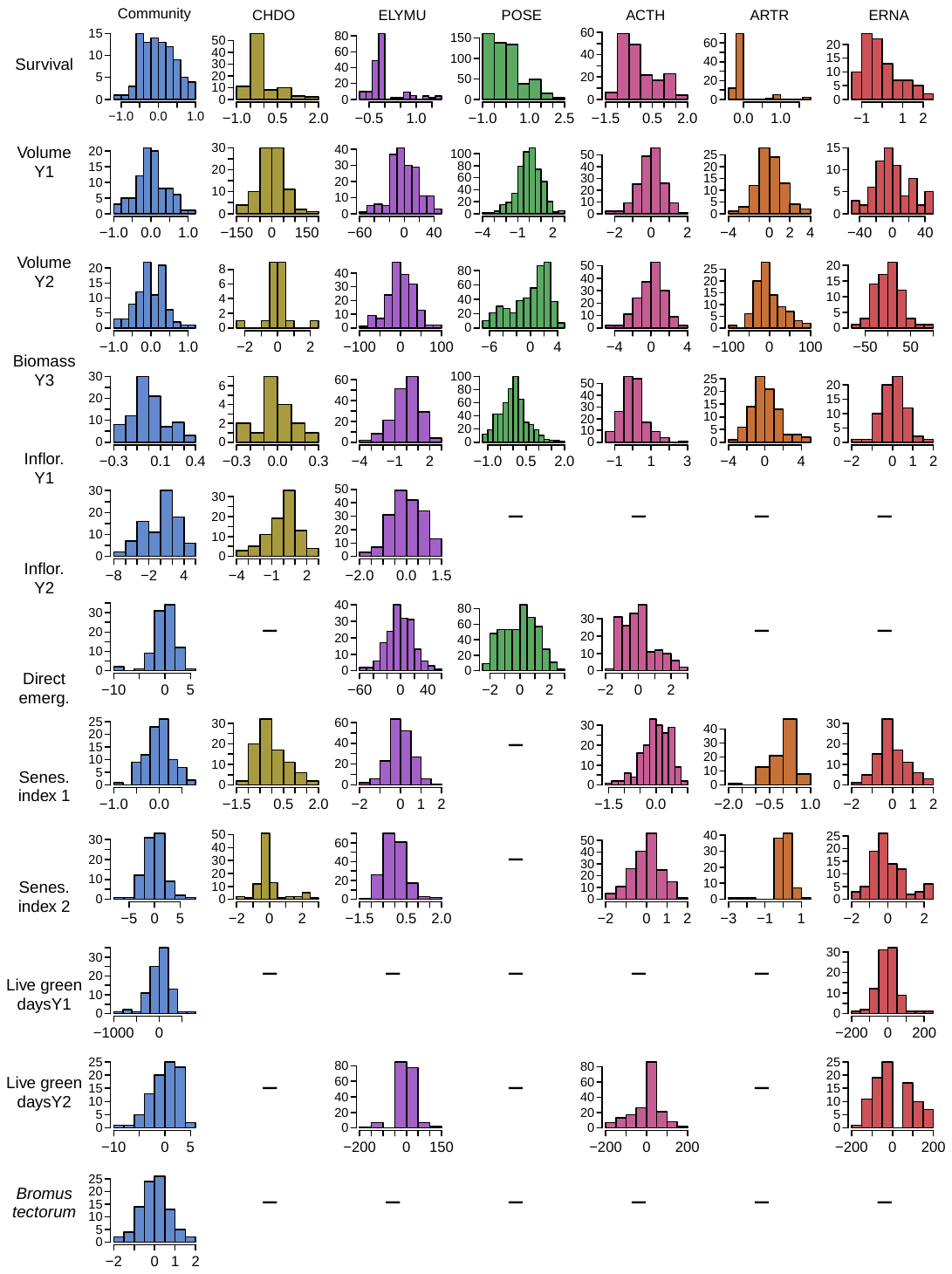
